# An epigenetic landscape governs early fate decision in cellular aging

**DOI:** 10.1101/630921

**Authors:** Yang Li, Yanfei Jiang, Julie Paxman, Richard O’Laughlin, Lorraine Pillus, Lev S. Tsimring, Jeff Hasty, Nan Hao

**Affiliations:** Section of Molecular Biology, Division of Biological Sciences, University of California San Diego, 9500 Gilman Drive, La Jolla, CA 92093, USA; Department of Bioengineering, University of California San Diego, La Jolla, CA 92093, USA; UCSD Moores Cancer Center, University of California San Diego, La Jolla, CA 92093, USA; BioCircuits Institute, University of California San Diego, La Jolla, CA 92093, USA

## Abstract

Chromatin instability and mitochondrial decline are conserved processes that contribute to cellular aging. Although both processes have been explored individually in the context of their distinct signaling pathways, the mechanism that determines *which* cell fate arises in isogenic cells is unknown. Here, we show that interactions between the chromatin silencing and mitochondrial pathways lead to an epigenetic landscape with multiple equilibrium states that represent different types of terminal cellular states. Interestingly, the structure of the landscape drives single-cell differentiation towards one of these states during aging, whereby the fate is determined quite early and is insensitive to intracellular noise. Guided by a quantitative model of the aging landscape, we genetically engineer a new “long-lived” equilibrium state that is characterized by a dramatically extended lifespan.

---

Cellular aging is a complex stochastic process. Many damage and stress factors contribute to aging, including chromatin instability, loss of proteostasis, mitochondrial dysfunction, reactive oxygen species, and others (*1, 2*). It has been generally considered that these factors accumulate during aging, resulting in cellular declines and eventually cell death. However, in each single cell, how individual aging factors conspire to drive the aging process remains unclear. For instance, cellular aging could be driven by multiple independent damage mechanisms that accumulate with varying rates, resulting in different aged phenotypes in different cells, or, alternatively, by the deterioration of the overall cellular condition, leading to a common aging and death pathway in all cells. A careful single-cell analysis is required to determine which of these scenarios actually underlies the cellular aging process. To this end, we investigated the replicative aging of yeast *S. cerevisiae*, a genetically tractable model for the aging of mitotic cell types such as stem cells (*3, 4*). Traditionally, yeast aging research has focused on “lifespan”, measured by the number of cell divisions before death, as a binary endpoint in defining the aging process (*5*). In this study, we integrated a newly-improved microfluidic device (Fig. S1) with time-lapse microscopy to measure lifespan along with dynamic changes of gene expression, chromatin state, and organelle morphology, enabling us to reveal a new “long-lived” mode of single-cell aging process.

Using single-cell imaging technologies, we found that isogenic WT cells exhibit two different types of phenotypic changes during aging (*6, 7*). About half of aging cells continuously produced daughters with a characteristic elongated morphology during later stages of lifespan. In contrast, the other half continuously produced small round daughter cells until death (Fig. 1A). We designate these two distinct modes of aging processes as “Mode 1” and “Mode 2”, respectively. Mode 2 features a more dramatic extension of cell cycle length and a shorter replicative lifespan than that of Mode 1 (Fig. S2).

**Fig. 1.**
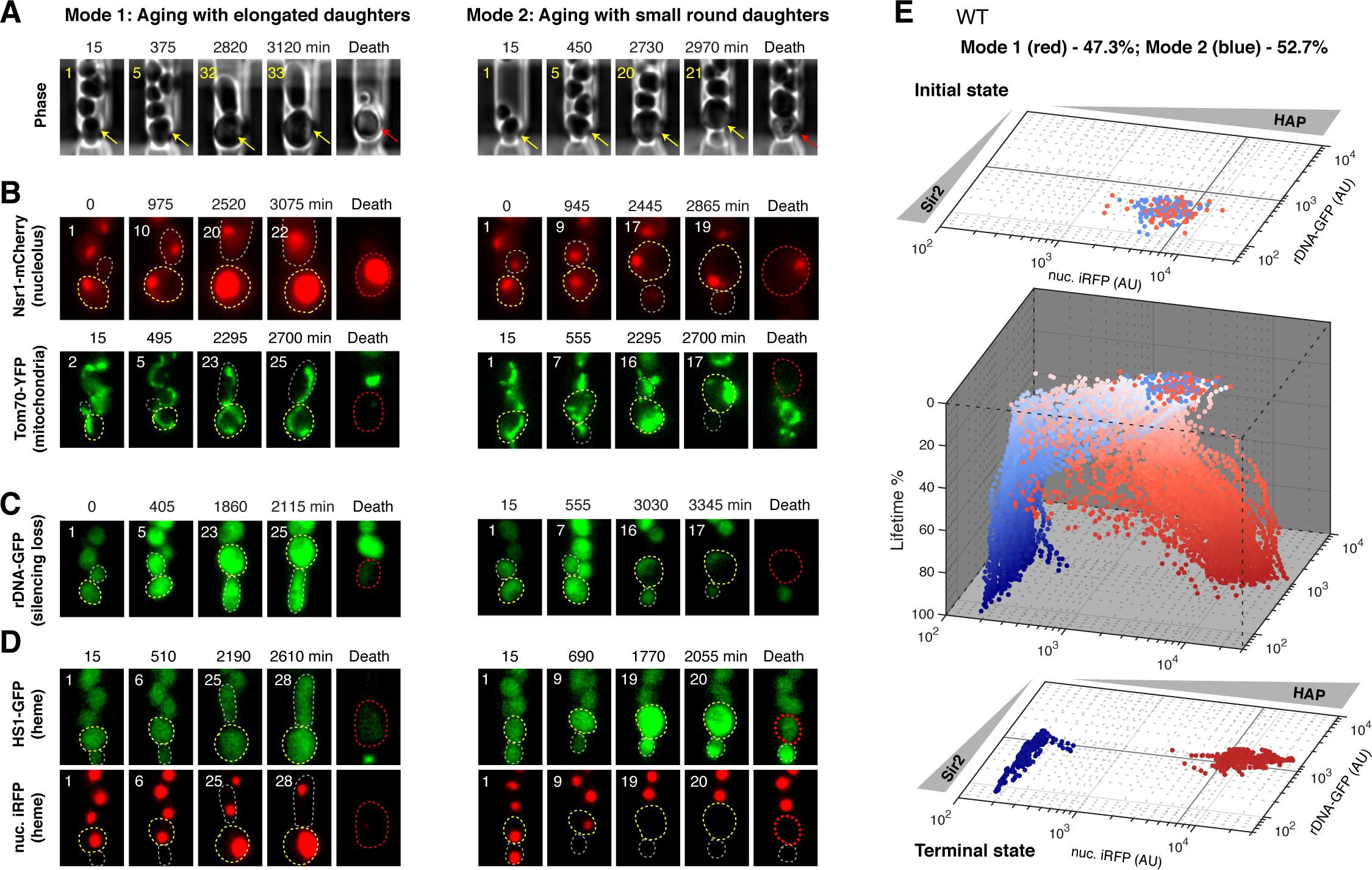
Sir2 and HAP mediate divergent aging of isogenic cells. Representative time-lapse images of Mode 1 and Mode 2 aging processes, for (A) phase (n=187), (B) Nsr1-mCherry (nucleolar marker; n=109) and Tom70-YFP (mitochondrial marker; n=142), (C) rDNA-GFP (n=256); and (D) HS1-GFP and nuc. iRFP (heme reporters in the same cells; n= 230). Time-lapse images are representatives of all Mode 1 and Mode 2 cells measured in this study. Replicative age of mother cell is shown at the top left corner of each image. For phase images, aging and dead mother cells are marked by yellow and red arrows, respectively. In fluorescence images, aging mother cells, newborn daughter cells, and dead mother cells are circled in yellow, grey and red, respectively. (E) Aging trajectories of a population of WT cells (n=187). Initial states of all the cells are projected onto a rDNA-GFP vs nuc. iRFP plane (reflecting Sir2 and HAP activities in single cells). The 3-dimentional space shows the aging processes of all the cells, in which z-axis represents the percentage of lifetime. Each dot represents quantified rDNA-GFP and nuc. iRFP fluorescence in a single cell at a given moment (red – Mode 1 cells; blue – Mode 2 cells). Terminal states of all the cells are projected onto another rDNA-GFP vs nuc. iRFP plane. Percentages of cells undergoing Mode 1 and Mode 2 aging are indicated at the top of the panel.

Aberrant structural changes of the nucleolus and mitochondria are major hallmarks of aging in yeast and many other organisms, indicating age-associated dysfunction of these organelles (*8, 9*). To determine the mechanistic basis of Mode 1 and Mode 2 aging, we tracked the morphologies of these organelles. We observed that the nucleoli were dramatically enlarged in all Mode 1 aged cells, but not in Mode 2 aged cells (Fig. 1B, top; Fig. S2C). Mitochondria in Mode 1 cells, however, retained a normal tubular morphology throughout the entire lifespan, whereas those in Mode 2 cells became aggregated before cell death (Fig. 1B, bottom). These results revealed that different modes of aging specifically target different organelles: Mode 1 aging leads to nucleolar decline, whereas Mode 2 aging results in mitochondrial decline.

Previous studies showed that age-dependent nucleolar enlargement is caused by instability of the ribosomal DNA (rDNA) (*10*), which is located within the nucleolus. A major mechanism of maintaining rDNA stability is chromatin silencing, which represses recombination of the highly repetitive rDNA. A conserved lysine deacetylase Sir2, encoded by the best-studied longevity gene to date, mediates rDNA silencing (*11*). We recently used a GFP reporter inserted at the non-transcribed spacer region of the rDNA (rDNA-GFP) to track rDNA silencing in single cells (*6*). Since the expression of the reporter is repressed by silencing, increased fluorescence indicates a loss of silencing. We found that Mode 1 cells, when aged, underwent sustained loss of rDNA silencing, whereas Mode 2 aged cells did not (Fig. 1C), in accord with nucleolar enlargement observed only in Mode 1 cells. Exposure to nicotinamide (NAM), an inhibitor of Sir2 that chemically represses its activity and disrupts rDNA silencing, induced a larger fraction (84.8%) of cells to undergo Mode 1 aging (Fig. S3A), confirming the causal role of Sir2 and rDNA silencing in driving Mode 1 aging. In addition, NAM treatment in liquid culture can also lead to the elongated cell morphology, confirming that this morphological change is a consequence of Sir2 activity loss (Fig. S3B).

For Mode 2 aging with mitochondrial decline, we considered the proposal that in human aging, deficiency of heme, a major functional form of iron, might be a key factor for mitochondrial decay (*12*). To determine how the heme level changes during aging, we used two independent reporters: a recently-developed fluorescent heme sensor (HS1)(*13*) and a nuclear anchored infra-red fluorescent protein reporter (nuc. iRFP)(*14*). For HS1, because binding of heme to the heme receptor quenches GFP fluorescence, an increased GFP signal indicates a decreased heme level. In contrast, for nuc. iRFP, since the fluorescence depends on biliverdin, a heme catabolism product, heme levels can be positively reflected by iRFP signals (Fig. S4, mechanisms and validation of heme reporters). These two reporters, co-expressed in the same cells, showed consistently that the heme level decreased sharply (within ∼300-600 minutes) in Mode 2 aging cells, but not in Mode 1 cells (Fig. 1D). Heme activates the heme activator protein (HAP) transcriptional complex to maintain mitochondrial biogenesis and function (*15*). Consistently, the expression levels of HAP-regulated genes, *COX5a* and *CYC1*, both of which encode key mitochondrial components, decreased specifically in Mode 2 aging cells (Fig. S5). Furthermore, when heme was artificially depleted through treatment with succinylacetone (SA), the majority (77.7%) of cells underwent Mode 2 aging (Fig. S3C), indicating that the decrease in heme and HAP level drives the Mode 2 aging process. Previous studies demonstrated that Pma1, a plasma membrane proton pump, increases during aging, causing vacuolar acidity decline and mitochondrial dysfunction (*9, 16, 17*). We found the age-dependent heme depletion converges at least partially with this Pma1-mediated aging pathway leading to mitochondrial decline (Fig. S6).

To determine how Sir2 and HAP operate together to drive single-cell aging, we introduced both rDNA-GFP and nuc. iRFP reporters into the same cells to simultaneously track the activities of Sir2 and HAP during aging. As shown in Fig. 1E, isogenic newborn cells began with relatively uniform levels of the two reporters. During aging, they diverged quite early in life (∼30% of lifetime) and progressed in two opposite directions. Mode 1 aging cells (47.3% of the population) ended life with high GFP and high iRFP levels, corresponding to a low Sir2, high HAP state. In contrast, Mode 2 cells (52.7%) ended at a high Sir2, low HAP state (Movie S1). These results revealed that genetically identical cells age toward two discrete terminal states with anti-correlated Sir2 and HAP activities, suggesting potential mutual inhibition between Sir2 and HAP that mediates the early-life divergence of aging processes.

To test this, we first deleted *SIR2* in the dual reporter strain. Most (83.1%) *sir2*Δ cells showed an elevated iRFP level during aging, indicating an increased HAP level in the absence of Sir2 (Fig. 2A). The expression of HAP-regulated genes, *COX5a* and *CYC1*, is also elevated in *sir2*Δ cells (Fig. S7), consistent with a recent RNA sequencing analysis showing that the transcript levels of many HAP-regulated genes are elevated in *sir2*Δ (*18*). As a result, the majority of *sir2*Δ cells aged with Mode 1 phenotypes and very quickly reached the low Sir2, high HAP state. When *HAP4*, encoding a major component of the HAP complex, was deleted, most (89.8%) *hap4*Δ cells showed decreased rDNA-GFP during aging, indicating elevated Sir2 activity and rDNA silencing in the absence of HAP (Fig. 2B). As a result, the majority of *hap4*Δ cells aged with Mode 2 phenotypes and approached the high Sir2, low HAP state before death. We further confirmed that HAP inhibits rDNA silencing through Sir2, as the inhibition is gone in *sir2*Δ (Fig. S8). Taken together, these results demonstrate that the mutual inhibitory interactions between Sir2 and HAP govern the fate decision of aging cells.

**Fig. 2.**
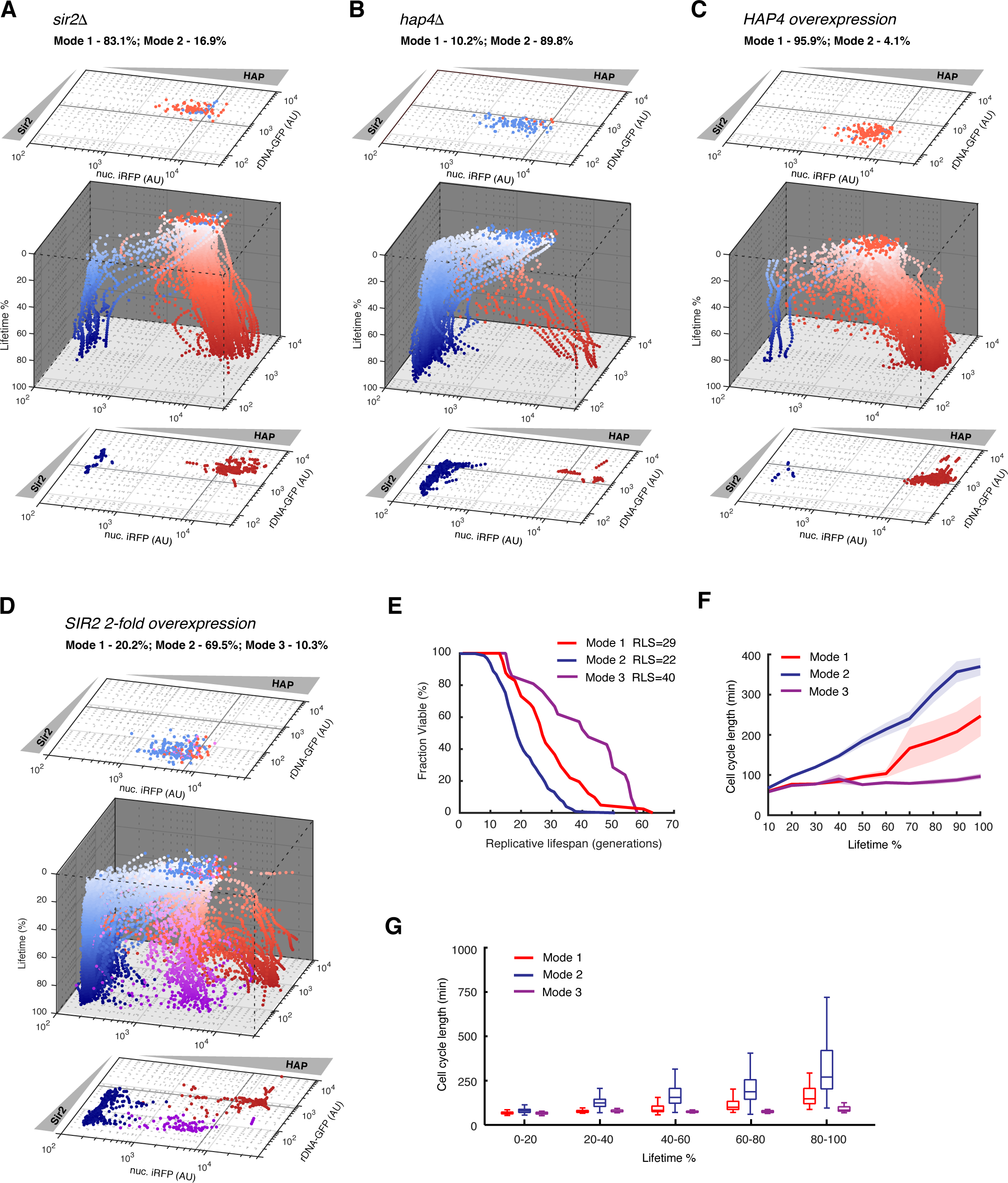
Genetic perturbations reveal mutual inhibitions between Sir2 and HAP and generate a new mode of aging. Aging trajecto-ries for (A) *sir2* Δ (n=89), (B) *hap4* Δ (n=98), (C) *HAP4* overexpression (n=118), and (D) *SIR2* 2-fold overexpression (n=203). Red – Mode 1 cells; blue – Mode 2 cells; purple – Mode 3 cells. (E) Replicative lifespans for Mode 1, 2 and 3 cells in the 2 × *SIR2* strain. (F) Changes of cell cycle length during Mode 1, 2 and 3 aging in the 2 × *SIR2* strain. The lifetime of each mother cell has been equally divided into 10 fractions. In each time fraction, the average length of all cell cycles has been quantified and plotted. Shaded areas represent standard errors of the mean (SEM). (G) Cell-to-cell variations in cell cycle length during Mode 1, 2, 3 aging in the 2 × SIR2 strain. The lifetime of each mother cell has been equally divided into 5 fractions. The distribution of all cell cycle lengths in each time fraction was plotted in the box plot. In the plot, the bottom and top of the box are first (the 25th percentile, q1) and third quartiles (the 75th percentile, q3); the band inside the box is the median; the whiskers cover the range between q1-1.5×(q3-q1) and q3+1.5×(q3-q1).

We next examined the effects of Hap4 and Sir2 overexpression, both of which can extend lifespan (*19, 20*). As expected from the deletion results, when Hap4 expression is increased, most (95.9%) cells showed decreased Sir2 activity and aged with Mode 1 phenotypes, confirming the inhibition of Sir2 by HAP during aging (Fig. 2C). In addition, caloric restriction (CR), which induces Hap4 expression (*21*), dramatically increased the proportion of Mode 1 aging cells (98% vs 52.7%) (Fig. S9), in accord with the results of Hap4 overexpression. In contrast, 2-fold overexpression of Sir2 increased the proportion of cells that underwent Mode 2 aging with decreased HAP activity, in agreement with Sir2-mediated inhibition of HAP. Intriguingly, however, in addition to Mode 1 and 2 aging, overexpression of Sir2 generated a third mode of aging that ended at a high Sir2, high HAP state (Fig. 2D). Strikingly, cells undergoing this new Mode 3 of aging were long-lived, with an average lifespan about twice of that of WT (Mode 3 RLS=40 vs WT RLS=21) (Fig. 2E). Although these cells produced elongated daughters as they aged, their cell cycle lengths retained unchanged throughout their entire lifespans, very different from that of Mode 1 cells (Fig. 2F and 2G; Movie S2).

To analyze the mechanisms that give rise to divergent aging and the emergence of this new aging mode, we devised a mathematical model based on our experimental observations. The model is composed of two stochastic ordinary differential equations incorporating the mutual inhibition between Sir2 and HAP as well as the positive auto-regulation of each (Fig. 3A). Sir2 deacetylates H4-Lys16, creating high-affinity nucleosome binding sites to promote further Sir2 recruitment. This positive feedback loop was proposed to account for the bistability in chromatin silencing (*22-24*). For HAP, heme induces transcription of Hap4 and activates the HAP complex, leading to increased expression of the tricarboxylic acid (TCA) cycle genes, which, in turn, increase the biosynthesis of heme, forming a positive feedback loop (*25*). Collectively, this network topology has the potential to generate multistability (*26*).

**Fig. 3.**
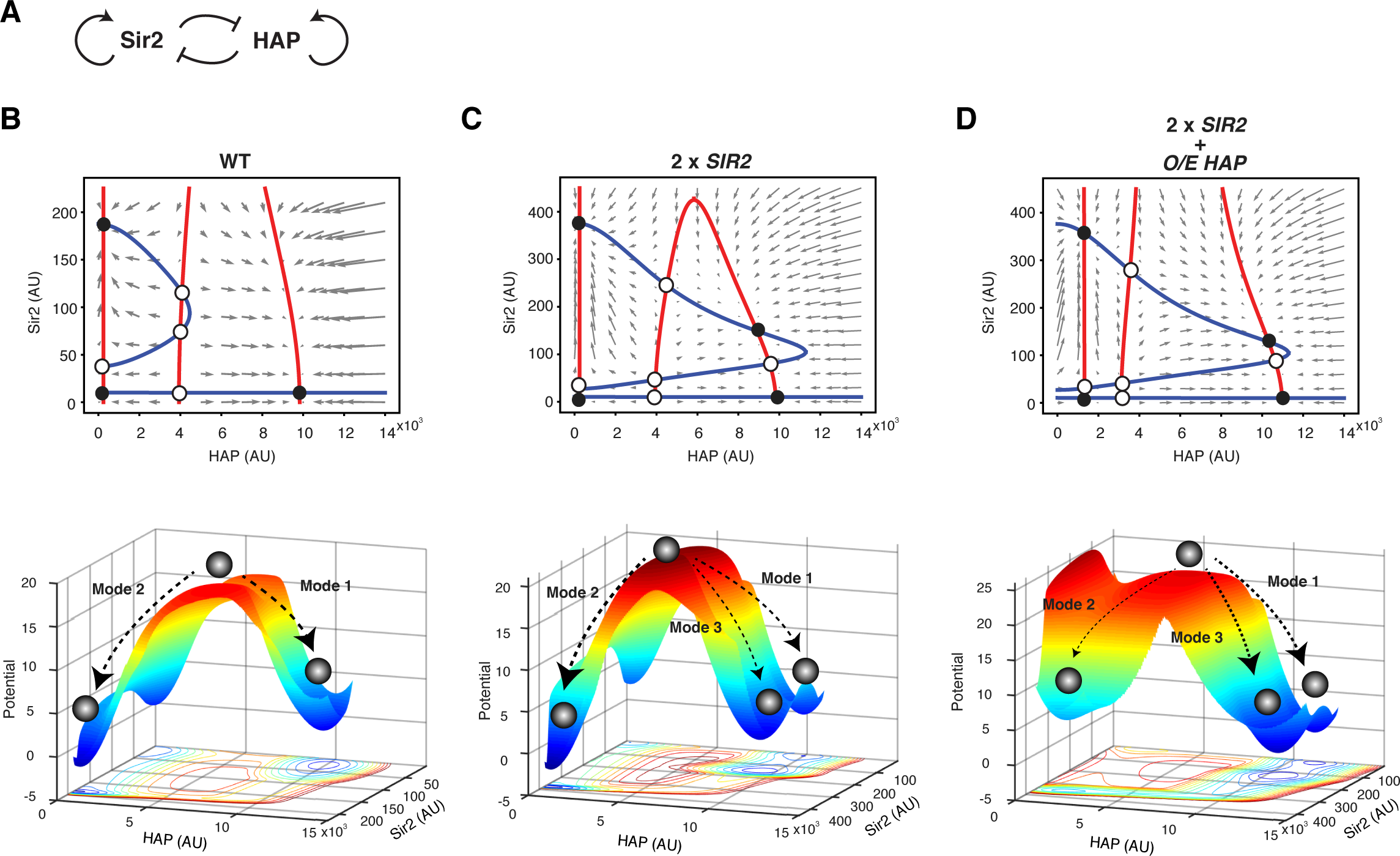
Computational modeling unravels multistability of the Sir2-HAP circuit that enables distinct modes of aging. (A) The diagram of the circuit topology. Graphic analyses of the model were conducted for (B) WT, (C) 2 × *SIR2*, and (D) 2 × *SIR2* + O/E *HAP* (overexpression). Top panels - phase plane diagrams. The nullclines for Sir2 and HAP are shown in blue and red, respectively. Grey arrows represent the vector field of the system, in which quivers show the rate and direction of movement. Fixed points are indicated with open (unstable) and filled (stable) circles. Bottom panels – computed potential landscapes. Well depths represent the probabilities of cells attracted to them. Single-cell aging processes can be depicted as a ball rolling into one of the wells on the surface.

The dynamical behavior of the system without noise can be analyzed graphically by plotting the nullclines and vector field in a Sir2-HAP phase plane. With appropriate parameters, there are seven fixed points for WT, among which three are stable (Fig. 3B, top). Although the positive feedback loops on Sir2 and HAP have the potential to enable quadrastability, the mutual inhibition with proper strength prevents their nullclines from intersecting in the high Sir2, high HAP region and hence eliminate one of the possible stable fixed points. To address the probabilistic nature of aging in single cells, we switched to a stochastic version of our model and computed an aging landscape by numerically solving the corresponding two-dimensional Fokker-Planck equation (Fig. 3B, bottom). Each of the stable fixed points of the deterministic system corresponds to a local well on the landscape, with the depth of the well reflecting its fate decision probability. Of the three wells, the two deep wells with low Sir2, high HAP and high Sir2, low HAP correspond to Mode 1 and Mode 2 aging processes observed experimentally. The low Sir2, low HAP well is shallower (reflecting lower probability) because the corresponding stable fixed point is close to an unstable fixed point. Some Mode 2 cells with relatively low Sir2 level (Fig. 1E) may belong to this stable fixed point, but their aging phenotypes appear to be the same as other Mode 2 cells with high Sir2, low HAP. (See Supplementary Information for the details about equations, parameters, bifurcation analysis, and landscape computation; Figs. S10-S14).

When the level of Sir2 is increased 2-fold, its increased activity can partially counteract the inhibition from HAP, leading to the emergence of a fourth fixed stable point on the phase plane and a new well on the potential landscape (Fig. 3C). The progression toward this new stable point with high Sir2 and high HAP corresponds to Mode 3 aging observed experimentally. However, at the same time, Sir2 overexpression biases the fate decision towards the high Sir2, low HAP state (deepest well on the landscape), in agreement with the experimental observation that a large fraction (69.5%) of cells underwent the short-lived Mode 2 aging (Fig. 2D). As a result, the average lifespan of the whole population was extended only modestly under this condition (RLS=26 vs WT RLS=21). We speculate that reducing the proportion of Mode 2 cells will further increase the lifespan of the whole population.

Based on our model, we predicted that an elevation in the HAP basal level can move the high Sir2, low HAP stable point and an unstable fixed point closer to each other, and hence reshape the landscape to bias towards the high HAP states, directing most cells to Mode 1 and Mode 3 aging and hence a longer lifespan (Fig. 3D). To test this prediction experimentally, we overexpressed both Sir2 and Hap4. Consistent with the model’s prediction, very few (5.8%) cells underwent Mode 2 aging and a large fraction (44.7%) of cells instead experienced long-lived Mode 3 (Fig. 4A). As a result, the combined perturbations effectively extended the lifespan of the population much more dramatically than overexpression of either Sir2 or Hap4 alone (Fig. 4B). It also enabled cells to maintain a relatively normal cell cycle length during aging (Fig. 4C), promoting healthy longevity of the whole population. From a network perspective, although overexpression of either Sir2 or Hap4 represses the other factor and pushes most cells to the other aging mode, combined perturbations of both can partially neutralize the mutual inhibition, potentiating a new long-lived aging mode with high Sir2 and HAP activities (Mode 3). Therefore, the perturbations of Sir2 and HAP, when combined, exhibit a synergistic, rather than additive, effect on lifespan extension (Fig. 4B). Our model could also help explain the synergy between CR (promoting HAP) and Sir2, two seemingly independent longevity factors, in a prior study (*27*).

**Fig. 4.**
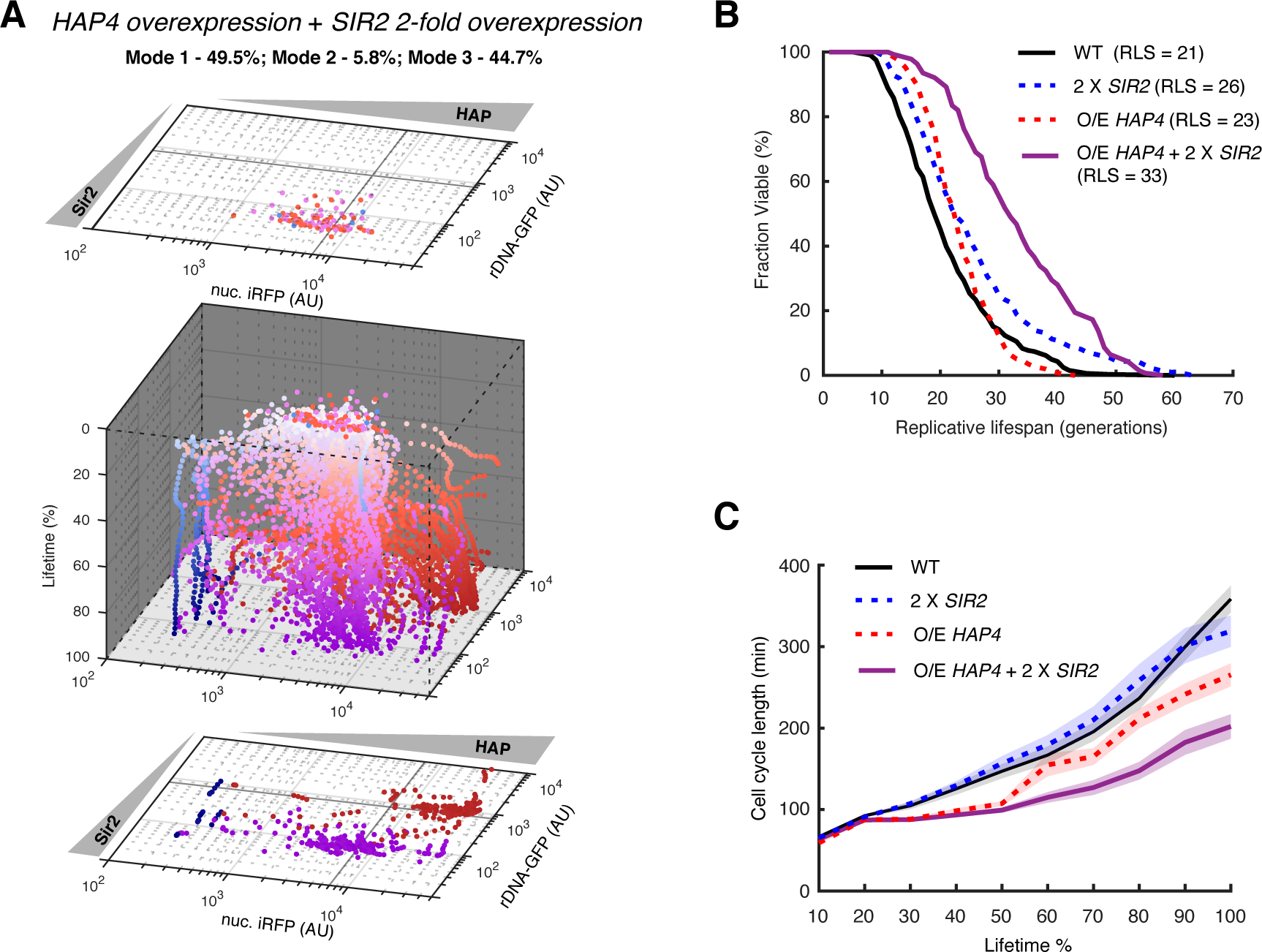
Combined perturbations of Sir2 and HAP reshape the aging landscape and synergistically promote longevity. (A) Aging trajectories for the strain with combined *HAP4* overexpression and *SIR2* 2-fold overexpression (n=103). Red – Mode 1 cells; blue – Mode 2 cells; purple – Mode 3 cells. (B) Replicative lifespans for WT, *SIR2* 2-fold overexpression (2 × *SIR2*), *HAP4* overexpression (O/E *HAP4*), and combined *HAP4* overexpression and *SIR2* 2-fold overexpression (O/E *HAP4* + 2 × *SIR2*). (C) Changes of cell cycle length during aging for WT, 2 × *SIR2*, O/E *HAP4*, and O/E *HAP4* + 2 × *SIR2*. Shaded areas represent SEM.

In summary, our results revealed that cellular aging can be considered as a programmable fate decision process rather than a consequence of passive damage accumulation. Individual cells initiate an aging-driven differentiation process quite early in life. They progress toward either silencing loss and nucleolar decline or heme depletion and mitochondrial decline, and age along the selected fate until death. Furthermore, based on the circuit dynamics, single-cell aging processes can be viewed as divergent progression towards one of the two wells on a Sir2-HAP landscape, corresponding to the low Sir2, high HAP and high Sir2, low HAP states. Importantly, model-guided perturbations of the circuit can reshape the aging landscape and enrich a long-lived mode of aging not observed in any WT cells, which leads to a dramatically extended lifespan of isogenic populations. This raises the possibility of rationally designing intervention strategies that target multiple network nodes, instead of single genes, to precisely modulate the aging dynamics of single cells or cell subpopulations and to more effectively promote healthy longevity.

## Acknowledgments

We thank G.Zhu and Y.Zhu for help with image analysis, and A.R.Reddi (Georgia Tech.) and D.E.Gottschling (Calico) for generously providing strains and reagents.

