## Supplementary Information for "An epigenetic landscape governs early fate decision in cellular aging"

### Supplementary Materials

- Figures S1-S9
- Computational Modeling + Figures S10-S14 + Tables S1-S2
- Materials and Methods + Tables S3-S4 + Figure S15
- Movies S1-S2

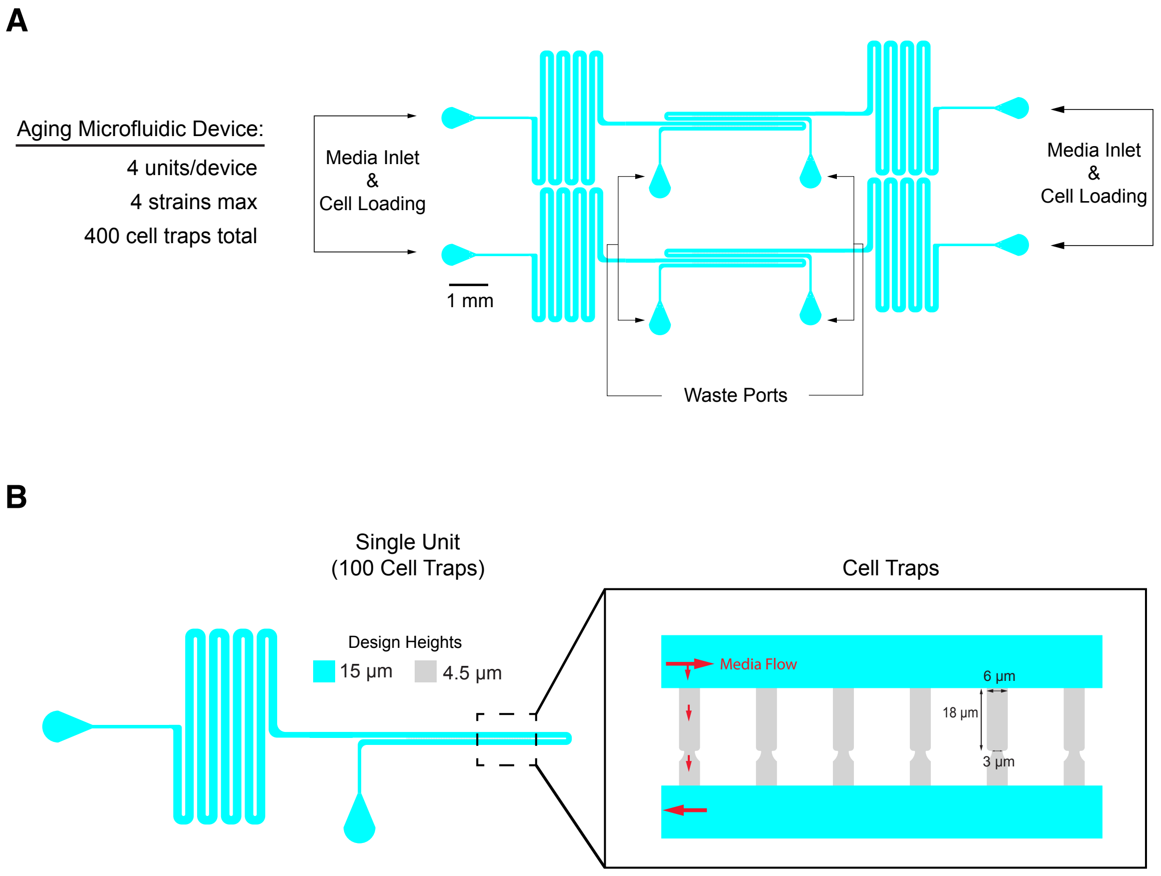


**Fig. S1. Microfluidic device for analyzing replicative aging in yeast.** (A) Overview of the device, which consists of four units with independent media inlets and waste ports. The device allows for a maximum of four different strains to be analyzed during a single experiment if necessary. Each device contains a total of 400 traps, with each individual unit having 100 cell traps. (B) Close-up of a single unit, consisting of two layers: media channels (blue, 15 μm tall) and cell traps (gray, 4.5 μm tall). Cells are loaded into chambers that trap mother cells while allowing them to bud continuously up toward the entrance of the trap or down through a small 3 μm opening toward the bottom of the trap.

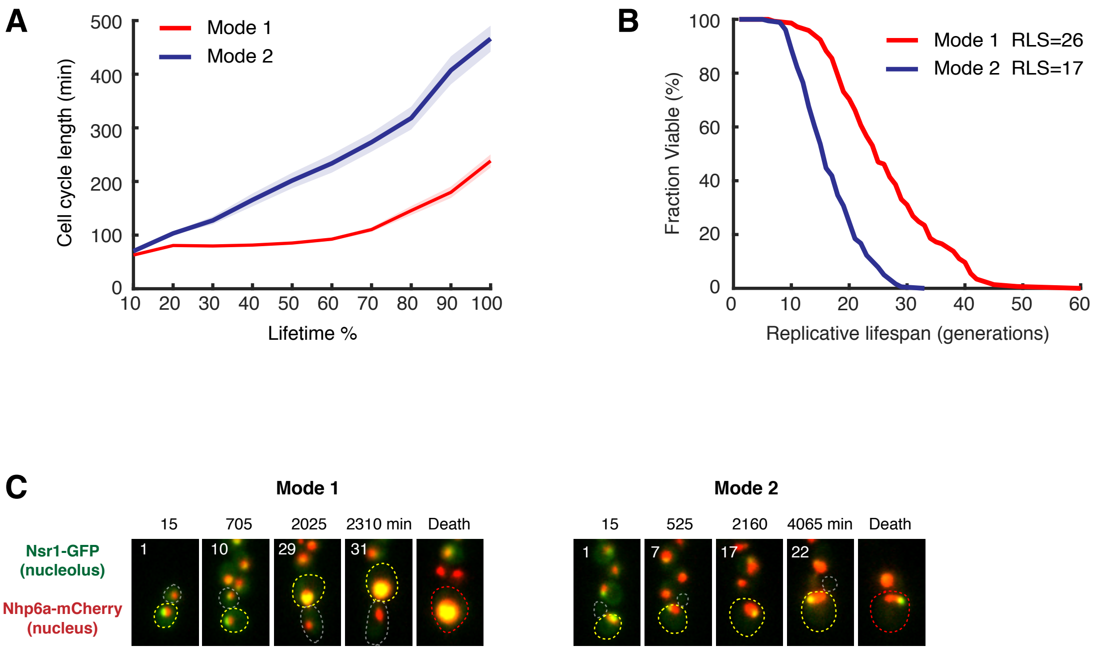


**Fig. S2. Mode 1 and Mode 2 aging show different cell cycle changes, lifespans, and nucleolar morphological changes.** (A) Changes of cell cycle length during Mode 1 (red) and Mode 2 (blue) aging (Mode 1: n=89; Mode 2: n=99). To quantifying the changes in cell cycle length during aging, the lifetime of each mother cell has been equally divided into ten fractions. In each fraction, the average length of all cell cycles of all mother cells has been quantified and plotted. Shaded areas represent standard errors of the mean (SEM). (B) Average replicative lifespans (RLSs) of Mode 1 and Mode 2 cells. (C) Distribution of the nucleolus within the nucleus during Mode 1 and Mode 2 aging. Representative time-lapse images of Nsr1-GFP (nucleolar marker; green) and Nph6a-mCherry (nuclear marker; red) during Mode 1 and Mode 2 aging processes were shown. Replicative age is shown at the top left corner of each image. Aging mother cells, newborn daughter cells, and dead mother cells are circled in yellow, grey and red, respectively. Ordinarily, yeast nucleoli are cresent-shaped structures confined to one edge of the nucleus. In Mode 1 aged cells, nucleoli were enlarged and occupied the entire nucleus, whereas those in Mode 2 aged cells remained the normal structure and size (n=119).

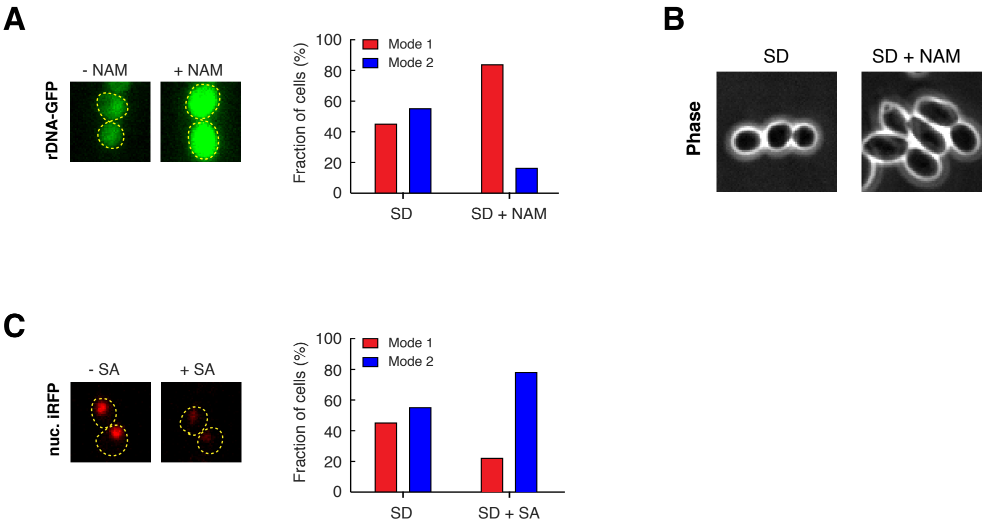


**Fig. S3. Chemical perturbations drive Mode 1 or Mode 2 aging.** (A) Nicotinamide (NAM) represses Sir2 and rDNA silencing, leading to an increased proportion of cells undergoing Mode 1 aging. Left: representative images illustrating the effects of NAM on the rDNA-GFP reporter; Right: proportions of Mode 1 (red) and Mode 2 (blue) aging cells in SD medium in the absence (n=187) or presence (n=138) of 5 mM NAM. (B) Yeast cells became elongated in liquid culture with SD + 5 mM NAM. Representative phase images of cells cultured in SD or SD + NAM for 10 hours are shown. These results confirmed that the elongated cell morphology is a phenotype of Sir2 activity loss, independent of microfluidic confinement. (C) Succinylacetone (SA) inhibits heme biosynthesis, leading to an increased proportion of cells undergoing Mode 2 aging. Left: representative images illustrating the effects of SA on the nuc. iRFP reporter; Right: proportions of Mode 1 (red) and Mode 2 (blue) aging cells in SD medium in the absence (n=187) or presence (n=94) of 0.05 mM SA.

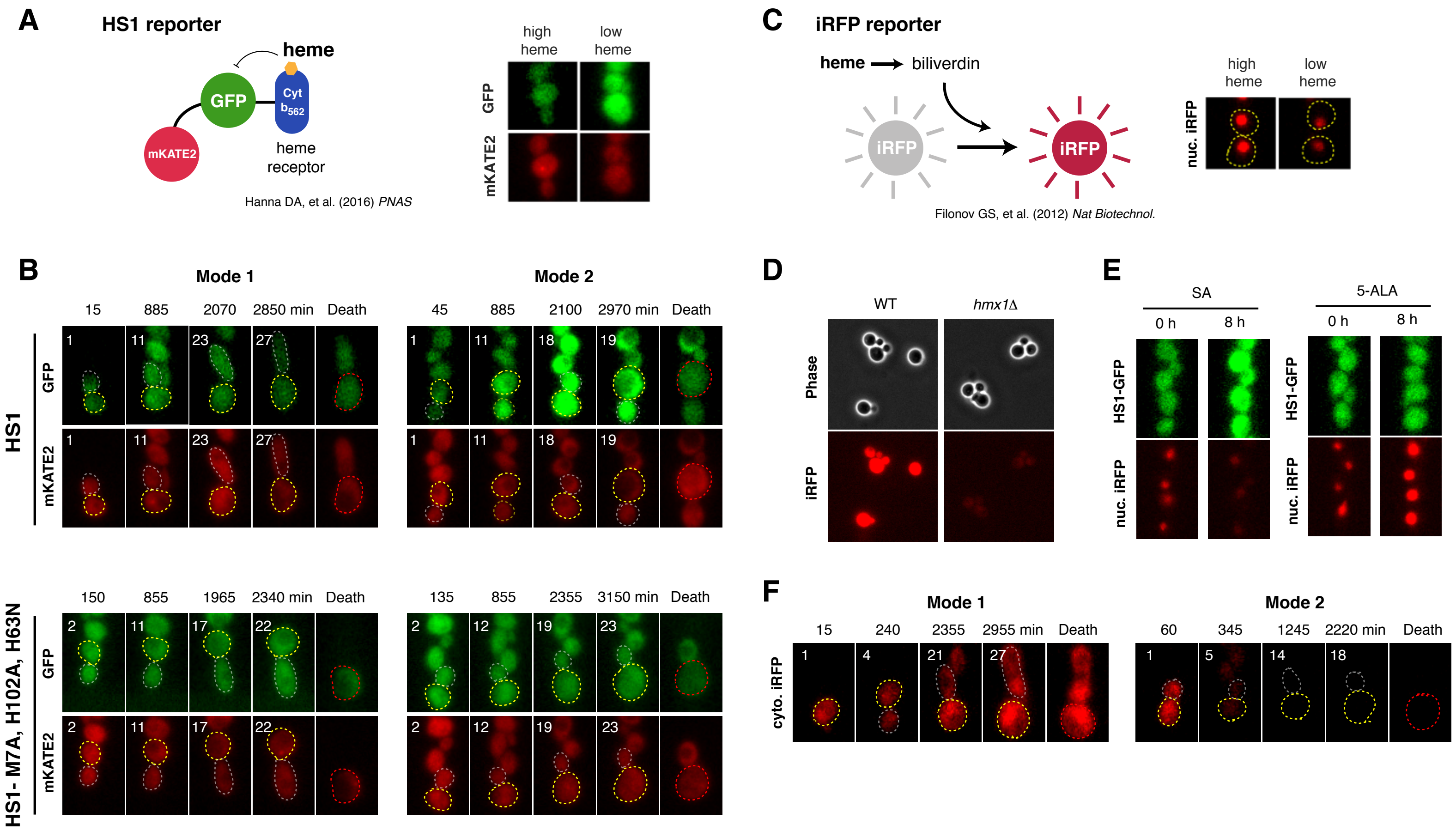


**Fig. S4. Validation of HS1 and iRFP reporters.** (A) Illustration of the ratiometric fluorescent heme sensor, HS1. Left: the reporter consists of a heme-binding domain (Cyt b562), fused to GFP and mKATE2. Right: because binding of heme to Cyt b562 quenches GFP fluorescence, but not mKATE2 fluorescence, a low GFP signal indicates a high heme level whereas a high GFP signal indicates a low heme level. mKATE2 is insensitive to the changes in heme level and serves as a control of reporter expression. (B) Responses of the HS1 reporter during Mode 1 and Mode 2 aging, with mKATE2 as an expression control. Top panel: representative time-lapse images of the HS1 responses during Mode 1 and Mode 2 aging processes. Replicative age is shown at the top left corner of each image. Aging mother cells, newborn daughter cells, and dead mother cells are circled in yellow, grey and red, respectively. For the HS1 reporter, both GFP and mKATE2 are shown. GFP fluorescence increased during Mode 2 aging, but not Mode 1 aging. mKATE2 fluorescence remained unchanged in both modes, confirming the expression of the reporter did not change in either mode of aging (n=230). Bottom panel: responses of a variant of HS1 with $His_{102}$ and $Met_7$ mutated to Ala and $His_{63}$ to Asn (HS1-M7A, H102A, H63N) during aging. This variant reporter is insensitive to heme level changes and therefore showed no changes in either mode of aging (n=183), confirming that the increase of HS1-GFP signal observed in Mode 2 aging is due to a change in heme level. These results also confirmed that the observed changes in rDNA-GFP and nuc. iRFP reporters during Mode 1 or Mode 2 aging (Fig. 1) are not caused by age-associated global effects on gene expression. (C) Illustration of the phytochrome-based near infra-red fluorescent protein (iRFP) reporter. Because the fluorescence of iRFP depends on biliverdin, a heme degradation product, high iRFP fluorescence indicates high heme level whereas low iRFP fluorescence indicates low heme level. (D) IRFP fluorescence in *hmx1*Δ cells. *HMX1* encodes the heme oxygenase that catalyzes the oxidative breakdown of heme to biliverdin. IRFP fluorescence was dramatically decreased in *hmx1*Δ cells, confirming the dependence of iRFP fluorescence on biliverdin. (E) Responses of HS1 and iRFP reporters to chemical perturbations of heme level. HS1 and iRFP reporters were co-expressed in the same cells. SA inhibits heme biosynthesis and decreases heme level. Consistently, increased HS1-GFP signal and decreased iRFP signal were observed in response to 1.5 mM SA for 8 hours. In contrast, increased iRFP signal was observed in response to 0.8 mM 5-aminolevulinic acid (5-ALA), a precursor in heme biosynthesis, which increases intracellular heme level. HS1, as a high-affinity heme sensor, is saturated and hence insensitive to the increase of heme level induced by 5-ALA. (F) Responses of cytoplasmic iRFP reporter during Mode 1 and Mode 2 aging (n=124). Cytoplasmic and nuclear-anchored (Fig. 1D) iRFP reporters showed similar patterns of fluorescence changes during aging, confirming that the iRFP signal changes during aging is independent of reporter localization. To facilitate single-cell tracking and image quantification, the nuclear-anchored iRFP reporter (nuc. iRFP) was used throughout this study.

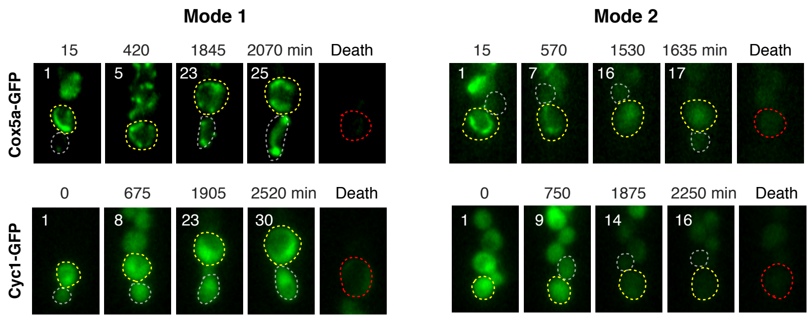


**Fig. S5. Expression levels of HAP-regulated genes in Mode 1 and Mode 2 aging.** Representative time-lapse images of Cox5a-GFP and Cyc1-GFP during Mode 1 and Mode 2 aging processes. Replicative age is shown at the top left corner of each image. Aging mother cells, newborn daughter cells, and dead mother cells are circled in yellow, grey and red, respectively. Levels of Cox5a-GFP (n=165) and Cyc1-GFP (n=179) decreased specifically during Mode 2 aging, but not Mode 1 aging.

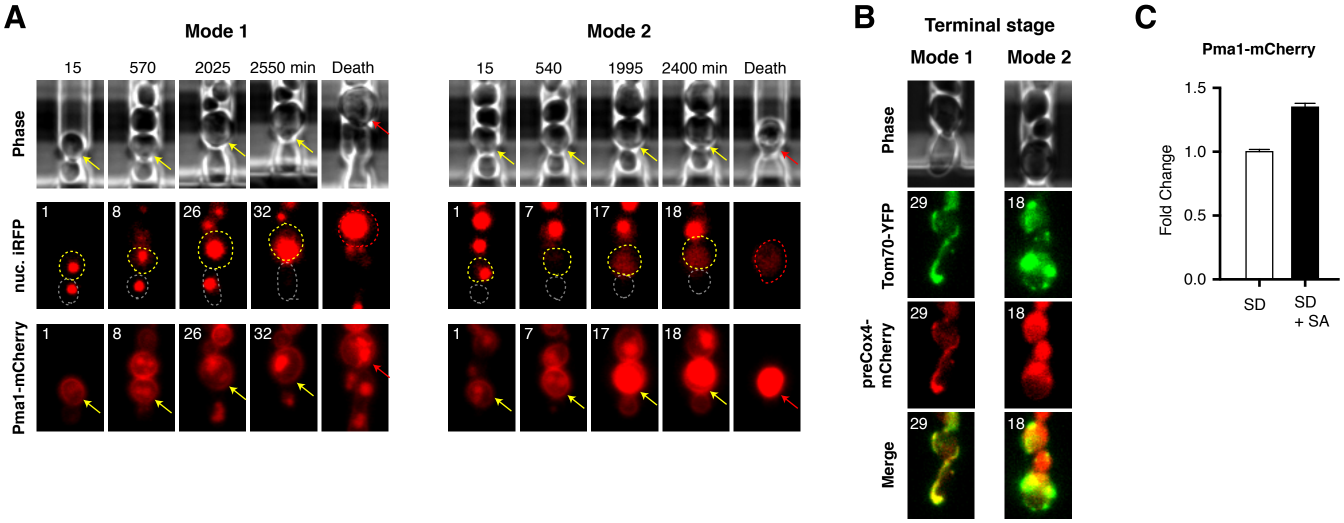


**Fig. S6. Convergence of age-dependent changes in heme and HAP onto the Pma1-mediated pathway leading to loss of mitochondrial membrane potential.** (A) Representative time-lapse images of Pma1-mCherry during Mode 1 and Mode 2 aging processes. Responses of the nuc. iRFP reporter from the same cells have also been included to reflect the convergence of changes in heme and Pma1 levels during Mode 2 aging. Replicative age is shown at the top left corner of each image. In the Pma1-mCherry images, aging mother cells and dead mother cells are denoted by yellow and red arrows, respectively. Previous studies showed that Pma1 is a proton ATPase that accumulates in aging cells and induces cytosolic pH increases, leading to reduced vacuolar acidity and loss of mitochondrial membrane potential. Concurrently with decreased heme level, Pma1 accumulated more dramatically during Mode 2 aging than Mode 1 aging (n=159), suggesting that the changes in heme level may converge onto the Pma1-mediated aging pathway. (B) Localization of proteins to the mitochondrial matrix in Mode 1 and Mode 2 aged cells. Representative images showed the localization of proteins to the outer mitochondrial membrane (Tom70-YFP) or the mitochondrial matrix (preCox4-mCherry) at the terminal stage of Mode 1 and Mode 2 aging (n=228). PreCox4-mCherry containing the inner mitochondrial membrane targeting pre-sequence of Cox4 fused with mCherry is a reporter for mitochondrial matrix targeting that depends on inner mitochondrial membrane potential. In Mode 1 aged cells, preCox4-mCherry remained colocalized to the tubular structure with Tom70-YFP; in contrast, in Mode 2 aged cells, preCox4-mCherry failed to localize to mitochondria and showed diffused cytoplasmic localization, indicating a loss of mitochondrial membrane potential. (C) Chemical depletion of heme induces the accumulation of Pma1-mCherry, suggesting a causal role of heme decline on Pma1 accumulation. Cells expressing Pma1-mCherry was treated with SA for 10 hours (n=116).

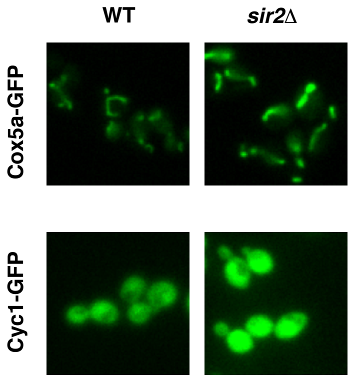


**Fig. S7. Deletion of Sir2 increases the expression levels of HAP-regulated genes.** Representative images showing Cox5a-GFP and Cyc1-GFP in WT and *sir2*Δ cells.

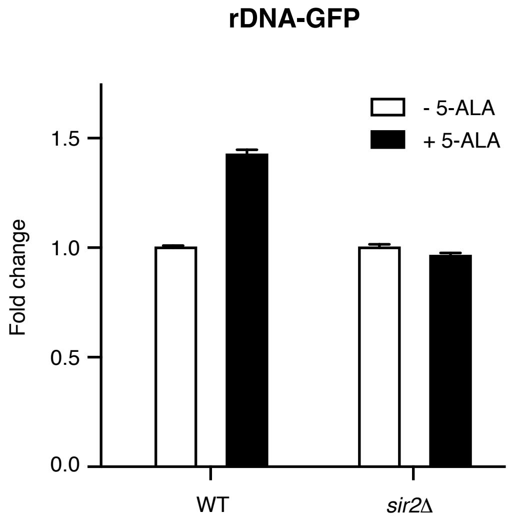


**Fig. S8. HAP inhibits rDNA silencing through Sir2.** The level of rDNA-GFP in WT cells increased in response to 10 hour treatment of 1.5 mM 5-aminolevulinic acid (5-ALA), a precursor in heme biosynthesis, which increases intracellular heme level. These results indicated that the elevated heme/HAP by 5-ALA decreases rDNA silencing (as reflected by increased rDNA-GFP). This inhibition is dependent on Sir2, since the effect of 5-ALA on rDNA silencing does not occur in the absence of Sir2.

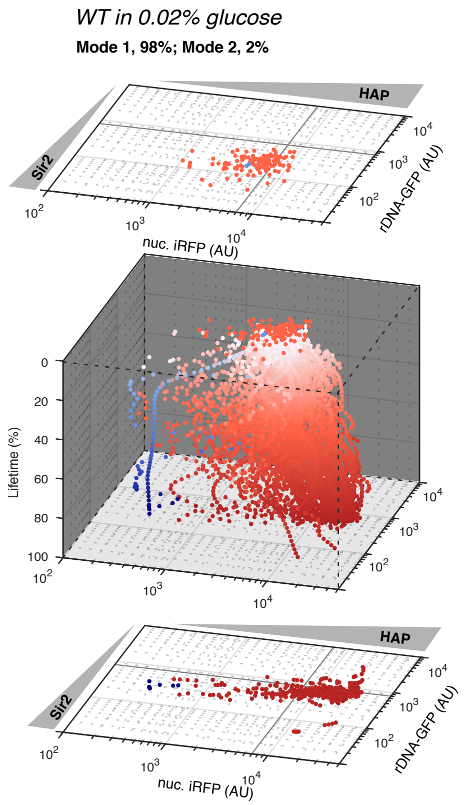


**Fig. S9. Aging trajectories under caloric restriction (CR, 0.02% glucose).** The vast majority (98%) of cells underwent Mode 1 aging with elongated daughters (red) and showed increasing rDNA-GFP and iRFP signals. Only 2% of cells underwent Mode 2 aging (blue). (n=138).

#### Computational Modeling

##### *Deterministic model*

In this model, we used two Hill functions of order 4 for both HAP and Sir2 positive feedbacks. The mutual inhibition was expressed by other two Hill functions of order 2 modulated by two strength factors, $\alpha$ and $\beta$. The difference between these two equations is that Sir2 (S) is regulated by activation and deactivation, while HAP (H) is regulated by synthesis and degradation. The nullclines were calculated numerically with the MATLAB "root" function and complex roots were omitted.

$$\frac{dH}{dt} = k_1 \frac{(1-\alpha)S^2 + K_{M1}^2}{S^2 + K_{M1}^2} \frac{H^4}{H^4 + K_{M2}^4} + k_2 - k_3 H \qquad (1)$$

$$\frac{dS}{dt} = k_4 \frac{(1-\beta)H^2 + K_{M3}^2}{H^2 + K_{M3}^2} \frac{S^4}{S^4 + K_{M4}^4} (S_{total} - S) + k_5 - k_6 S \qquad (2)$$

The parameters used in the model are listed below in Table S1.

**Table S1. Kinetic parameters used in the model for WT strain.**

| Parameter | Value | Description |
|---|---|---|
| $k_1$ | 40 $min^{-1}$ | HAP production rate |
| $k_2$ | 1 $min^{-1}$ | Basal level of HAP production rate |
| $k_3$ | 0.004 $min^{-1}$ | HAP degradation rate |
| $k_4$ | 0.02 $min^{-1}$ | Sir2 activation rate |
| $k_5$ | 0.04 $min^{-1}$ | Basal level of Sir2 activation rate |
| $k_6$ | 0.004 $min^{-1}$ | Sir2 deactivation rate |
| $K_{M1}$ | 280 | Equilibrium constant for Sir2 inhibition of HAP |
| $K_{M2}$ | 4250 | Equilibrium constant for HAP autoregulation |
| $K_{M3}$ | 2200 | Equilibrium constant for HAP inhibition of Sir2 |
| $K_{M4}$ | 90 | Equilibrium constant for Sir2 autoregulation |
| $\alpha$ | 0.35 | Strength factor for Sir2 inhibition of HAP |
| $\beta$ | 0.95 | Strength factor for HAP inhibition of Sir2 |
| $S_{total}$ | 225 | Total amount of Sir2 |

***Potential landscape computation***

Stochastic dynamics of aging can be described by the following Langevin equations in which two noise terms, $\xi_H$ and $\xi_S$, are added to the deterministic equations:

$$\frac{dH}{dt} = k_1 \frac{\left((1-\alpha)S^2 + K_{M1}^2\right)}{S^2 + K_{M1}^2} \frac{H^4}{H^4 + K_{M2}^4} + k_2 - k_3 H + \xi_H \quad (3)$$

$$\frac{dS}{dt} = k_4 \frac{\left((1-\beta)H^2 + K_{M3}^2\right)}{H^2 + K_{M3}^2} \frac{S^4}{S^4 + K_{M4}^4} (S_{total} - S) + k_5 - k_6 S + \xi_S \quad (4)$$

The noise terms are modeled as two independent white Gaussian processes with magnitudes $D_H, D_S$, $\langle \xi_i(t)\xi_j(t') \rangle = \sqrt{D_i\, D_j}\delta_{ij}\delta(t - t'); i, j \in \{H, S\}$. To generate potential landscapes $E(H, S)$, we deduced the associated two-dimensional Fokker-Planck equation from (3)(4) for the dynamics of the probability distribution $p(H, S, t)$,

$$\frac{\partial p(\mathrm{H,S,t})}{\partial \mathrm{t}} = -\frac{\partial}{\partial H}[F_H p(H, S, t)] - \frac{\partial}{\partial S}[F_S p(H, S, t)] + D_H \frac{\partial^2 p}{\partial H^2} + D_S \frac{\partial^2 p}{\partial S^2} \quad (5)$$

where $F_H, F_S$ are the right-hand sides of equations (1), (2). We simulated the Fokker-Planck equation in a rectangular domain $0 < H < H_{max}, 0 < S < S_{total}$ with no-flux boundary conditions (to conserve probability) until $t = T_{max}$ when the probability density converged to the asymptotic state $P(H, S)$ and used the formula $E(H, S) = -\log\big(P(H, S)\big)$ to obtain the potential landscape. As initial condition we used a Gaussian probability distribution $p(H, S, 0) = (2\pi\sigma_H\sigma_s)^{-1} exp\left[-\frac{(H-H_0)^2}{\sigma_H^2} - \frac{(S-S_0)^2}{\sigma_S^2}\right]$ (see Table S2 for the parameters of simulations). We employed first-order in time and second-order in space finite-difference scheme with 256 x 256 collocation points.

**Table S2. Parameters of numerical simulations of the Fokker-Planck model (WT strain).**

| | |
|---|---|
| $D_H$ | 500 |
| $D_S$ | 0.45 |
| $H_0$ | 3725 |
| $S_0$ | 112 |
| $\sigma_H$ | 5000 |
| $\sigma_s$ | 150 |
| $T_{max}$ | 5000 |
| $H_{max}$ | 15000 |
| Δt | 0.01 |
| ΔH | $H_{max}/256$ |
| ΔS | $S_{total}/256$ |

##### *HAP and Sir2 both have potential to generate bistability*

When α and β are set to be 0 in our model, i.e. the negative feedback terms $\frac{(1-\alpha)S^2+K_{M1}^2}{S^2+K_{M1}^2}$ and $\frac{(1-\beta)H^2+K_{M3}^2}{H^2+K_{M3}^2}$ are equal to 1, and the system does not have mutual inhibition. HAP and Sir2 form two independent positive feedback loops. These two positive feedbacks have 3 x 3=9 fixed points, four of which are stable. In the phase plane, the nullclines are intersecting vertical or horizontal lines (Fig. S10A). When α and β are set to non-zero values, the straight nullclines are distorted. In the WT case, α=0.35 and β=0.95, HAP has strong inhibition on Sir2. As a result, the Sir2 nullclines are greatly distorted, causing the disappearance of the $HAP_{High}$ $Sir2_{High}$ state (Fig. S10B).

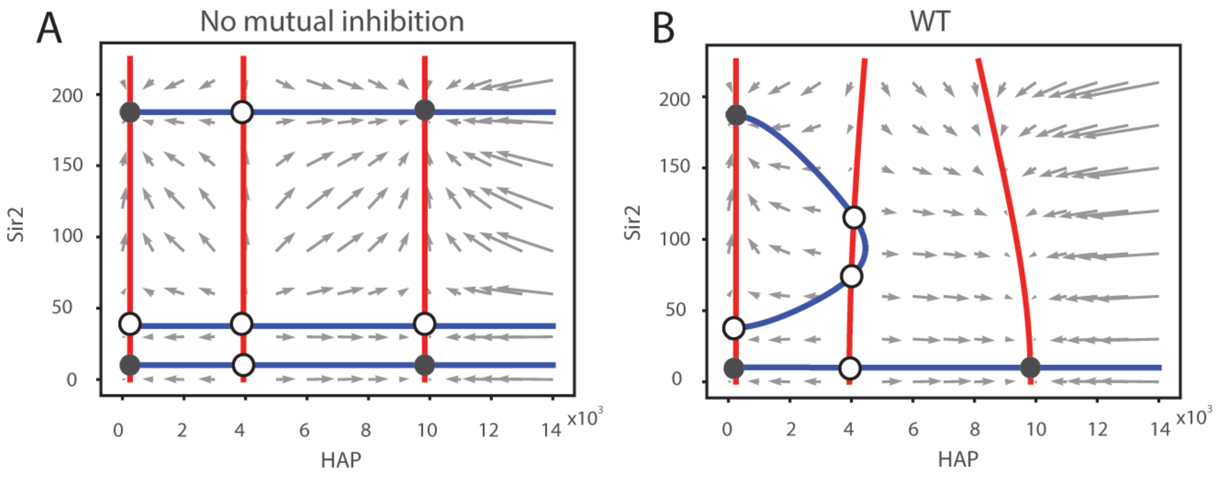


**Fig. S10. Phase planes and nullclines when the mutual inhibition is absent or present.** Red: HAP nullclines. Blue: Sir2 nullclines. (A) Phase plane in the absence of mutual inhibitions. Four stable fixed points are labeled with filled black circles, corresponding to the $HAP_{High}Sir2_{High}$, $HAP_{High}Sir2_{Low}$, $HAP_{Low}Sir2_{High}$, and $HAP_{Low}Sir2_{Low}$ states. Unstable fixed points are labeled with open circles. (B) Phase plane in the presence of mutual inhibitions. The $HAP_{High}Sir2_{High}$ state disappears when the nullclines are distorted by the mutual inhibition.

##### ***hap4*Δ and *sir2*Δ**

To simulate the aging process of the *sir2*Δ mutant, we set the total amount of Sir2 to a very low level (20 compared to 225 for WT; non-zero value to allow a comparison with the data using the rDNA-GFP reporter). In this case, Sir2 loses its bistability, as depicted by only one line (Fig. S11, blue) in the phase plane plot. This is consistent with the experimental results that *sir2*Δ mutant have similar extremely low Sir2 levels (as indicated by high rDNA-GFP levels) for both aging modes (Fig. 2A).

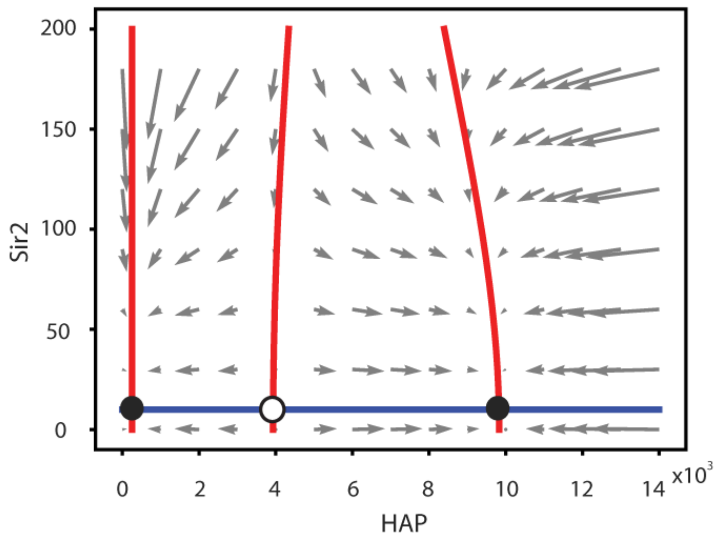


**Fig. S11. Phase plane and nullclines for the *sir2*Δ mutant.** Red: HAP nullclines. Blue: Sir2 nullclines.

For the *hap4*Δ mutant, as Hap4 is a component of the HAP complex, we assume the deletion of Hap4 does not completely abolish the HAP activity; instead, the mutant weakens the production of HAP ($k_1$ = 32 for this mutant). As shown in Fig. S12, the high HAP stable fixed point moves closer to the unstable fixed points, resulting in a bias fate decision toward Mode 2, in agreement with the experimental data of the *hap4*Δ mutant (Fig. 2B).

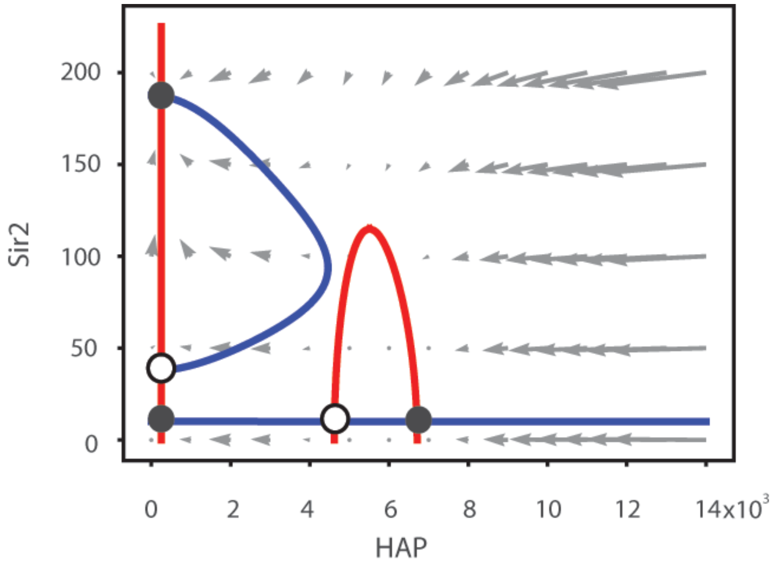


**Fig. S12. Phase plane and nullclines for the *hap4*Δ mutant.** Red: HAP nullclines. Blue: Sir2 nullclines.

##### *Dependence of the system's behaviors on total Sir2 level and the basal level of HAP production*

An increase in the total amount of Sir2 ($S_{total}$ = 450 instead of 225) can distort the HAP nullclines. More importantly, the increase in Sir2 can partially counteract the inhibition from HAP and lead to the emergence of the fourth stable point (Fig. 3C). The Sir2 nullclines are extended and intersect with the high HAP nullcline (compare Fig. S13, A and B). This new stable fixed point can account for the generation of Mode 3 in the 2 x *SIR2* strain observed experimentally (Fig. 2D). However, further increase in Sir2 (over 1200) makes the stable fixed point disappear again, due to the strong inhibition of Sir2 on HAP (lowered HAP nullclines in Fig. S13C). We plotted a bifurcation diagram to show that the Mode 3 aging happens within a critical range of Sir2 total amount, from 400 to 1200 (Fig. S13D).

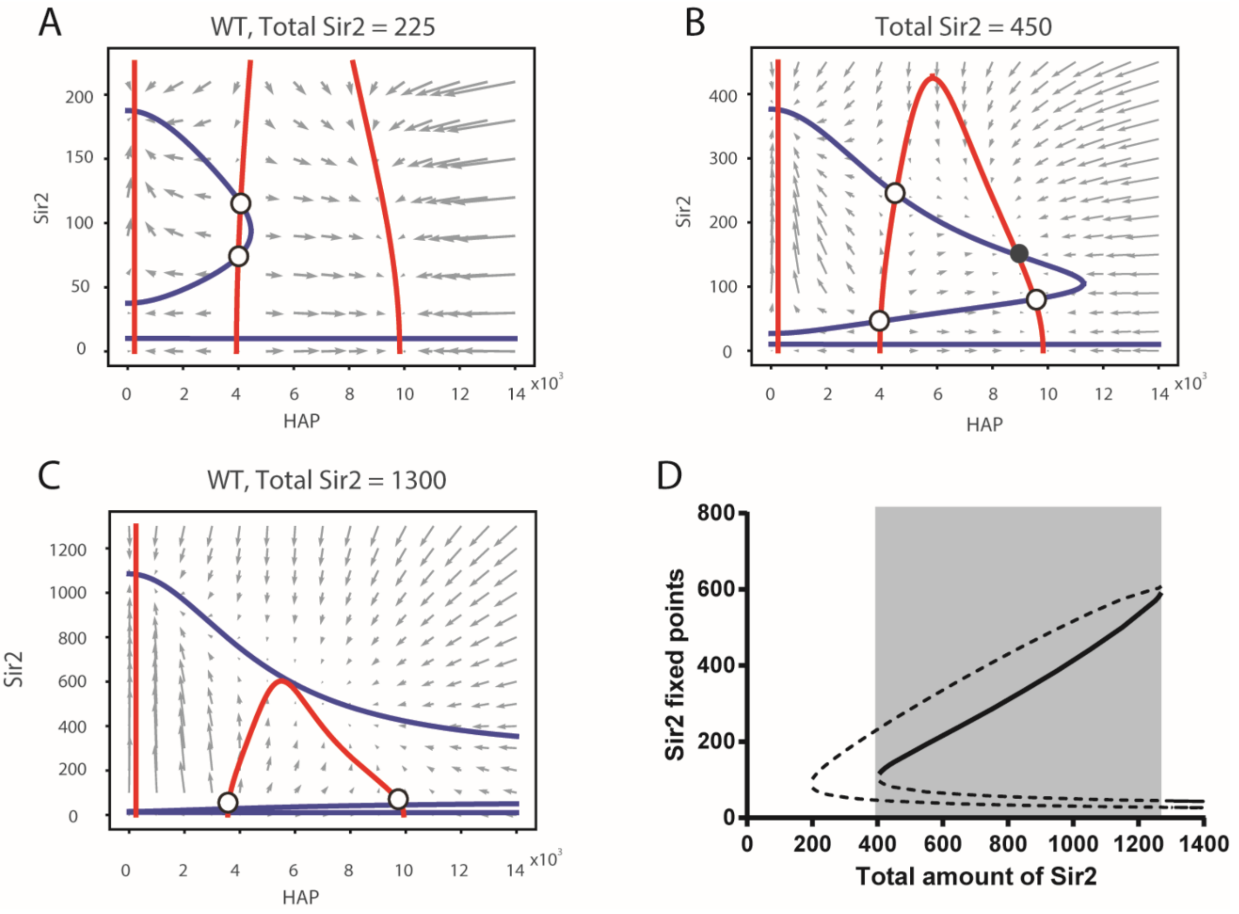


**Fig. S13. Dependence of the system's behavior on the total amount of Sir2.** (A-C) Phase planes for three different total Sir2 amounts. To generate the bifurcation diagram, only the fixed points that changes considerably (highlighted with circles, other fixed points are not highlighted) are plotted. (D) Bifurcation diagram for total amount of Sir2. Solid curve - stable fixed points. Dashed curve - unstable fixed points. The shaded area indicates the parameter range that enables the 4[th] stable fixed point (Mode 3 in the data).

We also investigated the stability dependence on the Hap4 basal expression ($k_2$). As shown in Fig. S14, the minimal amount of Sir2 needed to generate the new $Sir2_{high}HAP_{high}$ stable point is affected very modestly by $k_2$ (the lower bound of the shaded area). However, the upper limit of Sir2 greatly depends on the level of Hap4 production. This analysis led to the prediction that the combined overexpression of Sir2 and Hap4 can promote Mode 3 aging (Fig. 3D), which had been further tested experimentally (Fig. 4).

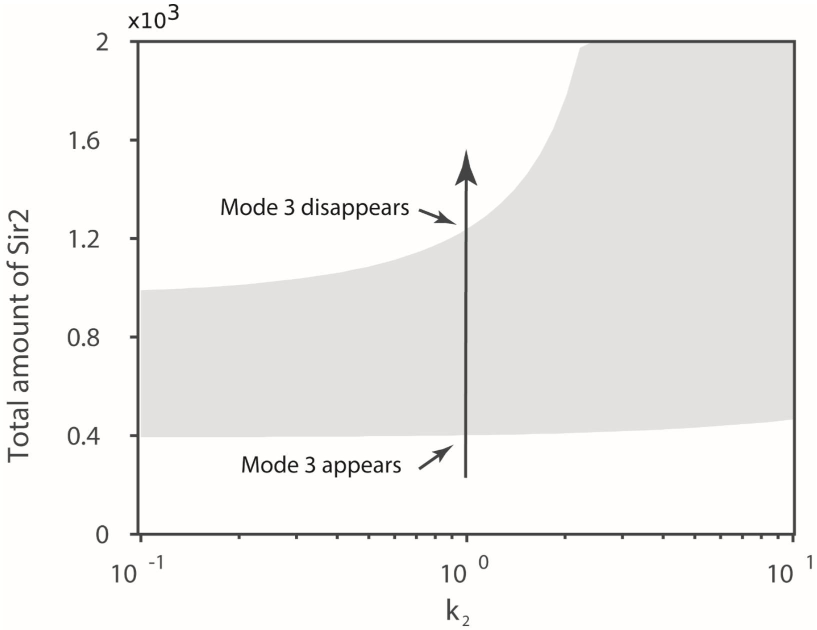


**Fig. S14. Stability dependence on the Hap4 basal expression (*$k_2$*).** The shaded area shows the region in which the $Sir2_{high}HAP_{high}$ stable fixed point (corresponding to Mode 3 aging) emerges in the parameter space. The long arrow corresponds to the changes of Sir2 amount in Figure S13D.

#### Materials and Methods

##### *Strains and plasmids construction*

Standard methods for the growth, maintenance and transformation of yeast and bacteria and for manipulation of DNA were used throughout. The yeast strains used in this study were generated from BY4741 (*MATa his3Δ1 leu2Δ0 met15Δ0 ura3Δ0*) strain background. Strain and plasmid information is provided in Tables S3 and S4.

To make the nuclear-anchored iRFP reporter, an iRFP*::KanMX* fragment was PCR amplified and integrated into the C-terminus of *NHP6a* at the native locus by homolog recombination. To make the *TOM70*-mCitrine(VA) reporter, a mCitrine(VA)*::HIS3* fragment was PCR amplified and integrated into the C-terminus of *TOM70* at the native locus by homologous recombination. Similarly, mCherry*::HIS3 or* GFP*::URA3* fragments were PCR amplified and integrated into the C-terminus of the target genes at its native locus by homologous recombination.

To make *sir2*Δ mutants, a *CgHIS3* fragment was amplified to replace the *SIR2* open reading frame by homolog recombination. Similarly, the *HAP4* ORF was replaced with *CgHIS3*. The *URA3* ORF is absent in the BY4741 background. To add a mutated *URA3* gene (*ura3-1*) back to its native locus, a *CgURA3* fragment was amplified and inserted at *URA3*, then *CgURA3* was replaced with a *ura3-1* allele from the W303 strain.

To make the *HS1* heme reporter (NHB0558), a *SalI_HS1_SacI* fragment from pDH013 (*13*) was ligated into the pRS306 vector. To make the $HS1^{M7A\ H102A,H63N}$ heme reporter (NHB0560), a *SalI_* $HS1^{M7A,\ H102A,H63N}$*_SacI* fragment from pJA010 (*13*) was ligated into the pRS306 vector. To make the cytosolic iRFP reporter, a *NotI_*$P_{TDH3}$*-iRFP-linker-GFP_SalI* fragment containing 680bp of the *TDH3* promoter, iRFP ORF and GFP ORF was made by fusion PCR and then ligated into pRS306, yielding plasmid NHB0454. *A SalI_*$P_{TDH3}$*-preCox4-mCherry_SacI* fragment was made by fusion PCR then ligated into pRS306 to get plasmid NHB0683. Plasmid NHB0658 and NHB0638 were constructed in a similar way.

Yeast strains with heme reporters were generated by transformation with either NHB0558 or NHB0560, which were digested with *StuI* for integration at *ura3-1*. The strain with cytosolic iRFP reporter was generated by transformation with NHB0454 digested with *ClaI* for integration at the promoter region of *NHP6a*. The strain with *preCOX4-mCherry* reporter (*17*) was generated by transformation with NHB0683, digested with *SnaBI* for integration at the promoter region of *TDH3*. The *HAP4* overexpression strain was generated by transformation with NHB0658, digested with *XbaI* for integration at the *HAP4* ORF. The strain with 2-fold overexpression of *SIR2* was generated by transformation with NHB0638, digested with *Sph1* for integration at the *SIR2* ORF. All transformations were performed with the standard lithium acetate method, and integration was confirmed by PCR.

We noted that BY4741 contains an in-frame Ty1 insertion near the 3' end of the *HAP1* gene. The resulting *hap1-Ty1* fused gene is expressed and replaces the 13 amino acids of *HAP1* C terminus with 32 amino acids from the Ty1 element (*28*). To exclude the possibility that the divergent aging and age-dependent heme decay revealed in this study are related to this mutation, we monitored phenotypic changes and nuc. iRFP dynamics during aging in the WT strain with the W303 genetic background (*MATa trp1 leu2 ura3 his3 can1 GAL+ psi+*), in which the *HAP1* gene is intact.

Similar to that observed in BY4741, about half of W303 cells underwent Mode 1 aging with elongated daughters and an increased heme level, whereas the other half underwent Mode 2 aging with a sharp decay in heme level (Fig. S15). These results confirmed that the divergent aging and age-dependent heme decay we observed are not specific to BY4741.

***Microfluidic device fabrication***
Design and fabrication of the microfluidic device for yeast replicative aging followed previously published work (*6*). In brief, SU8 2000 series photoresists (MicroChem), chrome glass masks (HTA Photomask) and an EVG620 contact mask aligner (EV Group) were used to construct and pattern the desired features for each layer onto silicon wafers (University Wafer Inc.). Feature heights were validated using a Dektak 150 surface profiler (Veeco).

***Setting up a microfluidic experiment***
Each chip contains four identical individual microfluidic devices. Each microfluidic device was checked carefully before use to ensure no dust or broken features were present. Before setting up a microfluidics experiment, the device was placed under vacuum for 20 min. After removing the device from the vacuum, all of the inlets of the device were immediately covered with 0.075% Tween 20 (Sigma- Aldrich Co.) for more than 5 min. The microfluidic device was placed on the stage of an inverted microscope with a 30°C incubator system. Media ports were connected to plastic tubing, which connected to 60 ml syringes with fresh SD (CSM powder from Sunrise Science, #1001-100) medium containing 0.04% Tween-20. The height the medium syringes is about 24 inches above the stage. The waste port of the microfluidic device was connected to plastic tubing, which was set to stage height. Yeast cells were inoculated into 2 ml of synthetic complete medium (SD, 2% dextrose) and cultured overnight at 30°C. 2 μl of saturated culture was diluted into 20 ml of fresh SD medium and grown at 30°C overnight until it reached $OD600_{nm} \sim 1.0$. For loading, cells were diluted by 4 - fold and transferred into a 60 ml syringe (Luer-Lok Tip, BD) connected to plastic tubing (TYGON, ID 0.020 IN, OD 0.060 IN, wall 0.020 IN). To load cells, the media port was replaced with a syringe filled with the yeast culture. The height of the cell loading syringe is also about 24 inches above the stage. The flow of medium in the device was maintained by gravity and drove cells into traps. Most traps were filled with cells within 1-2 minutes, after which the loading tubing was replaced with the media supply tubing and syringe. Heights of all tubing were adjusted to make the height difference around 60 inches. Waste medium was collected in a 50 ml tube to measure flow rate, which was about 2.5 ml/day. Note that Tween-20 is a non-ionic surfactant that helps reducing cell friction on the PDMS. We have validated previously that low concentrations of Tween-20 has no significant effect on cellular lifespan or physiology (*6*).

***Time-lapse microscopy***
Time-lapse microscopy experiments were performed using a Nikon Ti-E inverted fluorescence microscope with Perfect Focus, coupled with an EMCCD camera (Andor iXon X3 DU897). The light source is a spectra X LED system. Images were taken using a CFI plan Apochromat Lambda DM 60X oil immersion objective (NA 1.40 WD 0.13MM). During experiments, the microfluidic device was taped to a customized device holder inserted onto the motorized stage (with Encoders). In all experiments, the microscope was programmed to acquire images for each fluorescence channel every 15 min for a total of 80 hours or more. The exposure and intensity setting for each channel were set as follows: Phase 50 ms, GFP 10 ms at 10% lamp intensity with an EM Gain of

50, mCherry 50 ms at 10% lamp intensity with an EM Gain of 200, iRFP 300 ms at 15% lamp intensity with an EM Gain of 300, and YFP 100ms at 10% lamp intensity with an EM Gain of 300. The EM Gain settings are within the linear range. We had previously confirmed that this fluorescence imaging setting did not affect lifespan (*6*).

***Quantification of single-cell traces***

Fluorescence images were processed with a custom MATLAB code. Background of images from each channel were subtracted. Cell nuclei were identified by thresholding the iRFP images. Each image was evenly divided into 6 parts, each containing a single cell trap. The position of the dent in the cell trap was labeled. In each trap, the positions of the nuclei for single cells were labeled. Mother cells were identified by comparing the positions of the dent and the positions of the nuclei. For fluorescent reporters that are evenly diffused inside the cell, the nuclei of mother cells were further dilated to generate a mask to quantify the fluorescence intensities. The mean intensity value of the top 40% pixels of fluorescence reporter is quantified, as described previously (*6*). We have also tested segmenting the whole cell using phase images and quantified the mean fluorescence intensities of the whole cell. The resulting time traces were similar to those obtained using the nucleus-based segmentation. Because the nucleus-based segmentation is more robust and enables automated analysis of a large number of cells, we chose to use this method for all imaging analysis.

To plot the reporter level changes as a function of the percentage of lifetime, the total lifetime of each mother cell was equally divided into 50 fractions. The mean intensities of nuclear iRFP reporter and rDNA silencing reporter in each fraction were calculated and used for plotting.

Cell divisions of each mother cell were manually identified and counted at the time that the nuclei become completely separated between mother and daughter cells. To quantify the change of cell cycle length during aging, the lifetime of each single cell was equally divided into 10 fractions. The mean length of all cell cycles in each fraction was calculated and plotted.

***Categorization of aging modes***

Cells were categorized to different aging modes based on their aging phenotypes, characterized by the morphologies of their late daughters. Mothers continually producing round daughters at the last four generations were categorized as "Mode 2". Mothers continually producing elongated daughters at the last four generations were categorized as either "Mode 1" or "Mode 3". Among these cells, those that have an intensity of rDNA-GFP reporter below 200 (the cutoff obtained from the average initial state of the population) at the last cell cycle were categorized as "Mode 3", whereas cells that have an intensity of rDNA-GFP reporter above 200 (indicating low Sir2) at the last cell cycle were categorized as "Mode 1". A small fraction of cells showed abnormal morphologies even at the very beginning of the experiment and have a lifespan shorter than 5 generations. Those cells were excluded from analysis.

**Table S3. Strains used or constructed in this study.**

| Strain Name | Description |
|---|---|
| NH0270 | *BY4741 MAT*a *his3Δ1 leu2Δ0 met15Δ0 ura3Δ0, RDN1::NTS1-*$P_{TDH3}$*-GFP-URA3, NHP6a-iRFP-kanMX* |
| NH0277 | *BY4741 MAT*a *his3Δ1 leu2Δ0 met15Δ0 ura3Δ0, RDN1::NTS1-*$P_{TDH3}$*-GFP-URA3, NHP6a-iRFP-kanMX, sir2::CgHIS3* |
| NH0283 | *BY4741 MAT*a *his3Δ1 leu2Δ0 met15Δ0 ura3Δ0, NHP6a-iRFP-kanMX, ura3-1::URA3-*$P_{TDH3}$*-GFP* |
| NH0505 | *BY4741 MAT*a *his3Δ1 leu2Δ0 met15Δ0 ura3Δ0, RDN1::NTS1-*$P_{TDH3}$*-GFP-URA3, NHP6a-iRFP-kanMX, PMA1-mCherry-HIS3* |
| NH0599 | *BY4741 MAT*a *his3Δ1 leu2Δ0 met15Δ0 ura3Δ0, NHP6a-mCherry-HIS3,* $P_{NHP6a}$*::* $P_{NHP6a}$ *-iRFP-GFP-URA3* |
| NH0664 | *BY4741 MAT*a *his3Δ1 leu2Δ0 met15Δ0 ura3Δ0, NHP6a-iRFP-kanMX, ura3-1:: HS1-URA3* |
| NH0666 | *BY4741 MAT*a *his3Δ1 leu2Δ0 met15Δ0 ura3Δ0, NHP6a-iRFP-kanMX, ura3-1::* $HS1^{M7A, H102A,H63N}$*-URA3* |
| NH0717 | *BY4741 MAT*a *his3Δ1 leu2Δ0 met15Δ0 ura3Δ0, NHP6a-iRFP-kanMX, COX5a-GFP-URA3* |
| NH0719 | *BY4741 MAT*a *his3Δ1 leu2Δ0 met15Δ0 ura3Δ0, RDN1::NTS1-*$P_{TDH3}$*-GFP-URA3, NHP6a-iRFP-kanMX, hap4::CgHIS3* |
| NH0787 | *BY4741 MAT*a *his3Δ1 leu2Δ0 met15Δ0 ura3Δ0, NHP6a-iRFP-kanMX, CYC1-GFP-URA3* |
| NH0788 | *BY4741 MAT*a *his3Δ1 leu2Δ0 met15Δ0 ura3Δ0, NHP6a-iRFP-kanMX, HSP104-GFP-HIS3, NSR1-mCherry-URA3* |
| NH0804 | *BY4741 MAT*a *his3Δ1 leu2Δ0 met15Δ0 ura3Δ0, NHP6a-iRFP-kanMX, CYC1-GFP-URA3, sir2::CgHIS3* |
| NH0805 | *BY4741 MAT*a *his3Δ1 leu2Δ0 met15Δ0 ura3Δ0, NHP6a-iRFP-kanMX, COX5a-GFP-URA3, sir2::CgHIS3* |
| NH0868 | *BY4741 MAT*a *his3Δ1 leu2Δ0 met15Δ0 ura3Δ0, RDN1::NTS1-*$P_{TDH3}$*-GFP-URA3, NHP6a-iRFP-kanMX, HAP4::* $P_{TDH3}$*-HAP4-LEU2* |
| NH0880 | *BY4741 MAT*a *his3Δ1 leu2Δ0 met15Δ0 ura3Δ0, RDN1::NTS1-*$P_{TDH3}$*-GFP-URA3, NHP6a-iRFP-kanMX, HAP4::* $P_{TDH3}$*-HAP4-LEU2,* $P_{SIR2}$*::* $P_{SIR2}$ *-SIR2-HIS3* |
| NH0892 | *BY4741 MAT*a *his3Δ1 leu2Δ0 met15Δ0 ura3Δ0, NHP6a-iRFP-kanMX, TOM70-mCtrine(VA)-HIS3* |
| NH0897 | *BY4741 MAT*a *his3Δ1 leu2Δ0 met15Δ0 ura3Δ0, RDN1::NTS1-*$P_{TDH3}$*-GFP-URA3, NHP6a-iRFP-kanMX,* $P_{SIR2}$*::* $P_{SIR2}$ *-SIR2-HIS3* |
| NH0926 | *BY4741 MAT*a *his3Δ1 leu2Δ0 met15Δ0 ura3Δ0, NHP6a-iRFP-kanMX, TOM70-mCtrine(VA)-HIS,* $P_{TDH3}$*::* $P_{TDH3}$ *-preCOX4-mCherry-URA3* |
| NH0972 | *BY4741 MAT*a *his3Δ1 leu2Δ0 met15Δ0 ura3Δ0, NSR1-GFP-HIS3MX6, NHP6a-mCherry-LEU2* |

**Table S4. Plasmids constructed in this study.**

| Plasmid Name | Description |
| --- | --- |
| NHB0558 | pRS306-HS1 |
| NHB0560 | pRS306- $HS1^{M7A, H102A, H63N}$ |
| NHB0454 | PRS306-$P_{TDH3}$-iRFP-linker-GFP |
| NHB0658 | pRS305-$P_{TDH3}$-HAP4 |
| NHB0638 | pRS303-$P_{SIR2}$-SIR2 |
| NHB0683 | pRS306-$P_{TDH3}$-preCOX4-mCherry |

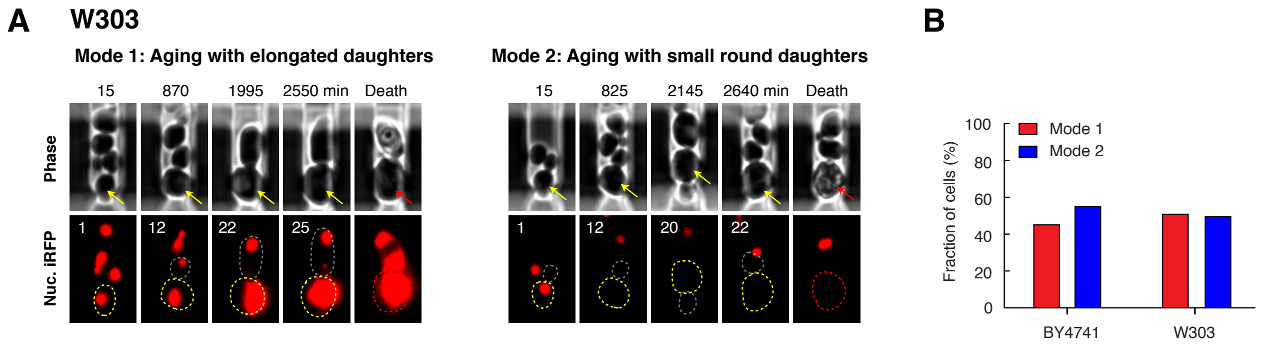


**Fig. S15. Divergent aging in the W303 WT strain.** (A) Representative time-lapse images of Mode 1 and Mode 2 aging processes in W303 WT, for phase (top) and nuc. iRFP (bottom). Time-lapse images are representatives of all Mode 1 and Mode 2 cells measured in this study (n=252). Replicative age is shown at the top left corner of each image. For phase images, aging and dead mother cells are denoted by yellow and red arrows, respectively. In fluorescence images, aging mother cells, newborn daughter cells, and dead mother cells are circled in yellow, grey and red, respectively. (B) Proportions of Mode 1 (red) and Mode 2 (blue) aging cells in BY4741 and W303. BY4741: Mode 1: n=89, Mode 2: n=99; W303: Mode 1: n=132, Mode 2: n=120. These results confirmed that the divergent aging and age-dependent changes of heme level revealed in this study are not specific to BY4741.

**Movie S1. Time-lapse movies showing fate decisions of isogenic WT cells during aging.** Right, movies of representative Mode 1 (top) and Mode 2 (bottom) aging cells (encircled) are played sequentially. RDNA-GFP reporter and nuc. iRFP reporter were co-expressed in the same cells and their fluorescence were measured during aging. Left, real-time quantification of reporter fluorescence plotted within a 3D aging space, in which z-axis represents the percentage of lifetime. After the aging trajectories of the two representative cells were quantified and plotted, trajectories of a population of isogenic WT cells were plotted in the space, generating Fig. 1E (Mode 1 – red; Mode 2 – blue).

**Movie S2. Time-lapse movies showing changes of cell cycle length during Mode 1, 2 and 3 aging in the 2 x *SIR2* strain.** Movies of representative Mode 1 (top), Mode 2 (middle), and Mode 3 (bottom) cells are played sequentially. Each movie contains time-lapse phase images of an aging mother cell (left) and real-time quantification of each cell cycle length (right), throughout its entire lifetime. The aging mother cell is denoted by an arrow. Red dots record the lengths of all cell cycles during the aging process of the mother cell (numbers on top of the red dots record the number of cell cycles – replicative age).
